# Trait-dependent diversification and spatio-ecological limits drive angiosperm diversity unevenness across the Canary Islands archipelago

**DOI:** 10.1101/2025.09.19.677265

**Authors:** Ryan F.A. Brewer, Ornela Dehayem Nanwou, Laura van Hoek, Marcelino José del Arco Aguilar, Yurena Arjona, William J. Baker, Noor M.S. van den Berg, Juli Caujapé-Castells, José María Fernández-Palacios, Cristina González-Montelongo, Ruth Jaén-Molina, Lucas S. Jansen, Louis S. Jay-García, Águedo Marrero, Sara Martín-Hernanz, Olivier Maurin, Raquel Negrão, Jairo Patiño, Stephan Scholz, Pablo Vargas, Alexandre R. Zuntini, Rampal S. Etienne, Luis Valente, Frederic Lens

## Abstract

Island biotas often show highly uneven species richness among lineages, influenced by clade age, diversification rates, and/or spatio-ecological limits. However, disentangling these drivers has been challenging due to the lack of comprehensive datasets across multiple lineages in the same geographical arena. The flora of the Canary Islands includes hundreds of plant lineages with contrasting species richness and harbours the highest number of species that evolved their woodiness in-situ (“insular woodiness”). Here, we present a phylogenomic reconstruction for Canary Island angiosperms and show that diversity unevenness in the flora is not driven by lineage age but by trait-dependent diversification and spatio-ecological limits. Our phylogenomic dataset, based on 1,244 newly generated and 501 published DNA sequences for 669 Canary Island species and 771 closely related mainland taxa, allows us to simultaneously study 435 plant lineages (∼50% of total). Applying dynamic stochastic modelling, we find the flora is shaped by high extinction and colonisation rates, maintaining a macroevolutionary equilibrium. Additionally, insular woody lineages exhibit higher diversification rates than the remaining flora. Our results suggest the uneven diversity of a highly dynamic insular region can be explained by the interaction of trait evolution and ecological constraints, providing insights into island biodiversity dynamics.

## Introduction

Explaining the macroevolutionary processes that have given rise to the unevenness in diversity across the tree of life is one of the major goals of (macro)ecology, evolutionary biology, and biogeography (1–3). Three main (non-mutually exclusive) explanations of this unevenness have been proposed (4): 1) variation in clade ages, with time differences being the sole explanatory factor (i.e., older clades have more species than younger clades; 5-7); 2) variation in diversification rates, emphasising clades accumulate species at differing rates through evolutionary time due to intrinsic characteristics of the clade (8,9); and 3) variable spatio-ecological limits on diversity over time (limits to species richness over time by resource limitations or the carrying capacity of a system/clade; 9-11). Underpinning these three hypotheses are the dynamic biogeographic processes of speciation, extinction, and colonisation, along with their interplay through time (12, 13). Integrating time-calibrated phylogenetic data with morphological and geographical information has become a powerful approach to better understand the mechanisms underlying these biodiversity patterns (14–18).

Island systems have long been at the forefront of evolutionary research (19–25). The relatively simple biotas of oceanic islands, their well-delimited boundaries compared to continental systems, and variations in the ages and diversities of colonising lineages simplify our interpretation of complex evolutionary processes (22, 23, 26). Island biogeography theory predicts that the processes of speciation, extinction, and immigration, as well as the age, area, and isolation of the island system, interact to explain how biodiversity accumulates in insular systems and tends towards an equilibrium (21, 29, 30). While the island equilibrium hypothesis has been contested (reviewed by 22, 31), a global test of MacArthur and Wilson’s predictions generally supports their conclusions (15). Nonetheless, analyses inferring the processes governing biodiversity patterns across the multiple lineages that make up an island assemblage are still in their infancy (15, 32–34).

Flowering plants (angiosperms) are one of the most diverse groups on islands, with insular systems harbouring over 60,000 endemic angiosperm species (25, 35). Studies on a global scale are beginning to disentangle how insularity affects the biogeographical processes driving angiosperm diversity (36). For example, certain traits or trait combinations are often markedly different in insular species than in their mainland relatives, a pattern known as ‘island syndrome’ (37–39). The most conspicuous island syndrome in angiosperms is insular woodiness (hereafter ‘IW’; 39-42), i.e. the tendency for herbaceous colonisers to evolve woodiness following island colonisation. Carine et al. (43) posit that the shift towards IW increases diversification rate (speciation minus extinction) in Macaronesian lineages, which was corroborated by other studies on the Canary Islands (41, 42). In contrast, Zizka et al (39) found that 51% of IW lineages globally contain only one to two species, challenging this IW-diversification hypothesis. Yet, to date, empirical tests of the effect of IW on diversification are restricted to single clades (44).

To investigate the processes governing insular biodiversity patterns, we selected the iconic angiosperm flora of the Canary Islands. The biota of this oceanic archipelago is one of the best studied globally and has a long history of evolutionary research (43, 45–47). Moreover, the Canarian angiosperm flora has one of the highest levels of species richness of any oceanic island archipelago (>1,300 species), harbours marked unevenness with numerous singleton lineages (composed of a single species) and many radiations (lineages with multiple species resulting from in-situ diversification), and has the highest documented number of shifts towards IW globally (27, 35, 39, 42, 48). Despite the potential of the Canarian flora as a model to study diversity unevenness across lineages, a comprehensive standardised phylogenetic dataset that allows for simultaneously analysing multiple Canary Island angiosperm lineages to test for causes of this unevenness under the same approach is lacking. In fact, such a dataset does not currently exist for any insular flora in the world.

Here, we analyse the processes that shaped Canary Island angiosperm biodiversity patterns in three ways. First, to quantify the number of independent colonisation events, the number of radiations and species per lineage, and how these vary amongst families across all native angiosperm genera of the archipelago, we update the estimated number of independent colonisation events using the phylogenetics literature and new phylogenetic data generated in this study. Second, to estimate time of island colonisation and speciation for Canary Island lineages, we provide a comprehensive standardised time-calibrated phylogenomics dataset comprising 669 species of non-monocot angiosperms native to the Canary Islands and their mainland relatives, using hybrid capture target enrichment of the Angiosperms353 bait set (49) and 166 fossil calibration points across angiosperms (50). Monocots were excluded from our target group for the phylogenetic tree because we are interested in the effects of IW on diversification, and monocots cannot produce wood or evolve woodiness (41, 51). To produce this phylogenomic dataset, we combined our newly generated sequences with existing published sequences of more distantly related mainland species (both monocots and non-monocots) to account for all the internal nodes that are associated with the fossils used for calibration (52). From this unprecedented phylogenomic dataset, we were able to estimate the time of island colonisation for hundreds of Canary Island plant lineages, allowing us to statistically assess the relationship between lineage age and diversity for ∼50% of the native angiosperm flora and ∼60% of the native Canary non-monocot species. Third, to estimate colonisation and diversification rates, compare rates of diversification between IW lineages and other lineages, and infer spatio-ecological limits to diversity, we fitted a dynamic stochastic model of island biogeography (DAISIE, Dynamic Assembly of Island biota through Speciation, Immigration, and Extinction; 14) to the time-calibrated phylogenetic information. DAISIE allows us to estimate lineage-specific rates of speciation, extinction, and colonisation for the complete Canary Islands non-monocot angiosperm assemblage, as well as clade-level diversity carrying capacity (14, 15).

By combining these three approaches, we provide an archipelago-wide empirical assessment of evolutionary rates of the Canary Islands. This allows us to test whether the unevenness in species diversity between lineages can be best explained by the clade age, diversification rate or spatio-ecological limits hypothesis, and to investigate how insular woodiness affects diversification rates on the archipelago.

## Results and Discussion

### The Canary Island angiosperm flora is characterised by a high number of colonisations

Our updated list of the native Canary Islands angiosperm flora based on published checklists recognises 1,385 species (608 endemics, ∼44%), representing 427 genera from 83 families, or 1,320 species across 400 genera when removing native but possibly introduced species (Supplementary Data S1, S2; 53-55). Combining this list with phylogenetic data, we identify 881 ‘likely’ colonisation events of the Canary Islands (minimum: 772, maximum: 968; Supplementary Data S1, see Methods), with the vast majority of colonisations being non-monocot angiosperms (676 likely colonisations, 605 minimum, 752 maximum; Supplementary Data S1). Flowering plant diversity is deeply uneven between lineages, ranging from one to 63 species per Canarian lineage. At least 480 lineages are singletons, 27 have more than five species, and six have 20 or more species (Supplementary Data S1). Colonisation of the Canary Islands is widespread across the angiosperm tree of life, spanning 33 angiosperm orders (Figure 1, Supplementary Data S1). The families with the highest numbers of colonisations are Asteraceae (249 species, 125 likely colonisations), Poaceae (130 species, 121 likely colonisations), and Fabaceae (130 species, 85 likely colonisations) (Figures 1, 2). Native insular species richness per family is positively correlated with the number of colonisations (Figure 1; Supplementary Data S3; Kendall’s rank correlation test, τ = 0.741, *p* < 0.01), with eight of the ten most species-rich families overlapping with the ten families with the highest number of colonisations (Supplementary Data S3). Native insular species richness per family is also positively correlated with mainland species richness (τ = 0.544, *p* < 0.01), where we use the Mediterranean Basin as our estimated mainland source pool, as this region accounts for 74 of the 83 families native to the Canaries (see Methodology). The most diverse mainland families also have the most colonisations of the Canaries (τ = 0.505, *p* < 0.01). Among the ten most species-rich families on the Canaries, only Crassulaceae (whose Canarian diversity is mostly the product of the largest radiation on the archipelago) is neither in the ten most frequently colonising families, nor in the ten most diverse mainland families, reflecting the importance of in-situ diversification in this family. Conversely, families commonly assumed to have a strong dispersal ability have very few native Canarian species per lineage but high native species richness (i.e. resulting from multiple colonisations). For example, Poaceae (130 species; 121 colonisations; 1.07 average native Canarian species per lineage), Cyperaceae (27/27/1), and Amaranthaceae (23/22/1.05) are represented almost entirely by single-species lineages (singletons) but are in the 15 most species-rich families (Supplementary Data S3).

**Figure 1.**
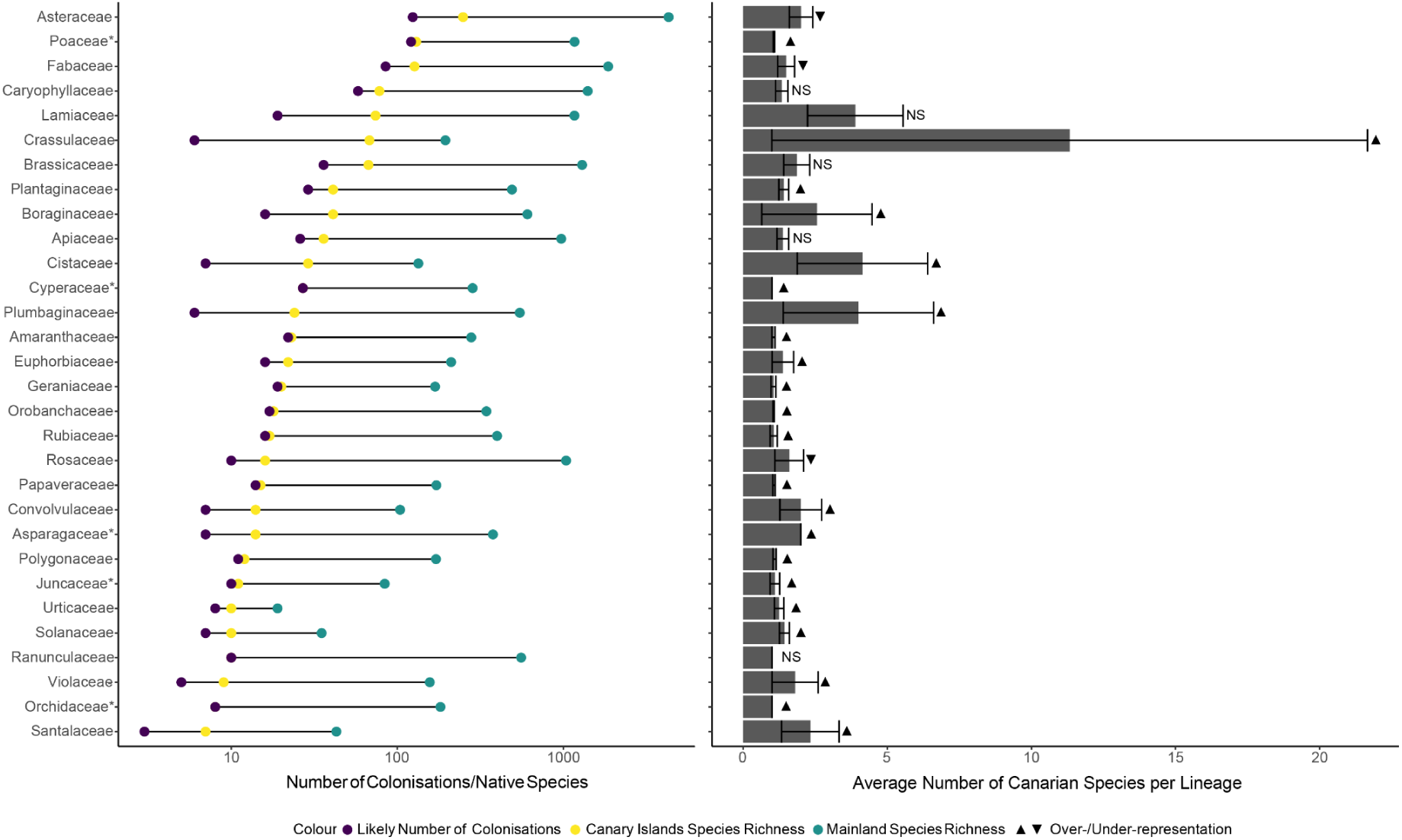
Top 30 most species-rich angiosperm families on the Canary Islands. a) Number of colonisations, total Canary Island species richness and mainland richness per family. b) Average number of native Canarian species (both endemic and non-endemic) per lineage and over-/under-representation relative to the mainland, where error bars indicate standard error. Significant island disharmony in comparison to the mainland is represented by triangles, upwards for over-represented, downwards for under-represented, and NS for not significant disharmony. Families with an asterisk are non-monocot angiosperms. Values for all families native to the Canaries are given in Supplementary Data S3 and S4.

Our phylogenetically informed estimate of 881 angiosperm colonisations of the Canary Islands (Supplementary Data S1, S2) is over 50% higher than previous estimates for all vascular plants. Using a stricter definition of native insular diversity, only considering species that are ‘Surely Native’ (54), Price et al. (47) estimated 479 colonisations of native vascular plants. When restricting our results to only include ‘Surely Native’ species from (54), we estimate 282 angiosperm colonisations (Supplementary Table S1). The drastic difference between the estimated number of likely colonisations and colonisations of ‘Surely Native’ species reflects the ‘Probably Native’ and ‘Possibly Native’ species included in our study. Nevertheless, our estimate of in-situ diversification (44%, Supplementary Data S1) is similar to the values obtained by Price et al. (47) and Schrader et al. (25), at 47% and 42%, respectively.

Using binomial tests to assess island disharmony, we identify 22 of the top 30 species-rich angiosperm families as significantly overrepresented on the Canaries relative to the mainland Mediterranean Basin source pool, with only three families (Asteraceae, Fabaceae, and Rosaceae) being underrepresented (Figure 1). Interestingly, these three families are the only ones among the 74 families in total that are under-represented on the Canaries compared to the Mediterranean mainland, a finding that is in line with recent studies on island disharmony that found Asteraceae, Fabaceae, and Rosaceae are consistently underrepresented on islands globally (25, 35, 56). Of the remaining 71 families native to the Canaries present in the Mediterranean Basin source pool, 58 are over-represented and 13 are equally represented (Supplementary Data S4).

### Time-calibrated phylogenetic framework of Canary Island angiosperms

To estimate colonisation and branching times across the Canary Islands angiosperm flora, we built a time-calibrated phylogenomic tree of the Canary Islands angiosperms, focusing on the non-monocot flora. We focus on non-monocot angiosperms for sampling and evolutionary analyses as monocots never produce wood, and to not bias our analyses against the effects of IW on diversification. Our phylogenetic reconstruction is based on 669 native Canarian species (∼60% of native Canarian non-monocot angiosperms, ∼50% of all native angiosperms) across 683 tips, and 771 closely related mainland taxa of Canarian species that are used as outgroups for each Canarian lineage. In total, we sequenced 1,244 new samples for this study, with the remaining tips from previously published studies (57–59), adding another 210 tips. We embedded these Canarian lineages into a broader phylogenetic angiosperm framework containing an additional 291 tips (thus adding up to a total of 1,745 tips; (52)), to enable reconstructing a time-calibrated evolutionary history across all sampled Canarian lineages using the same phylogenetic framework and fossil calibrations (see ‘Sampling Strategy’ in Methods; Figure 2, Supplementary Data S2, S5, S6, S7). While our focus is on non-monocots, we do include existing non-Canary Islands monocot sequence data in our tree (59 of the 501 additional tips), but only to assist with time calibration (to take advantage of the many existing monocot fossils). For phylogenetic support values (Felsenstein’s bootstrap proportions) for each lineage, see Supplementary Data S8. The 669 native Canarian species and 771 mainland relatives sampled in our tree represent 435 of our 676 estimated non-monocot lineages (∼65%) and 186 of the 241 IW Canarian species (77%; Figure 2; Supplementary Data S9).

**Figure 2.**
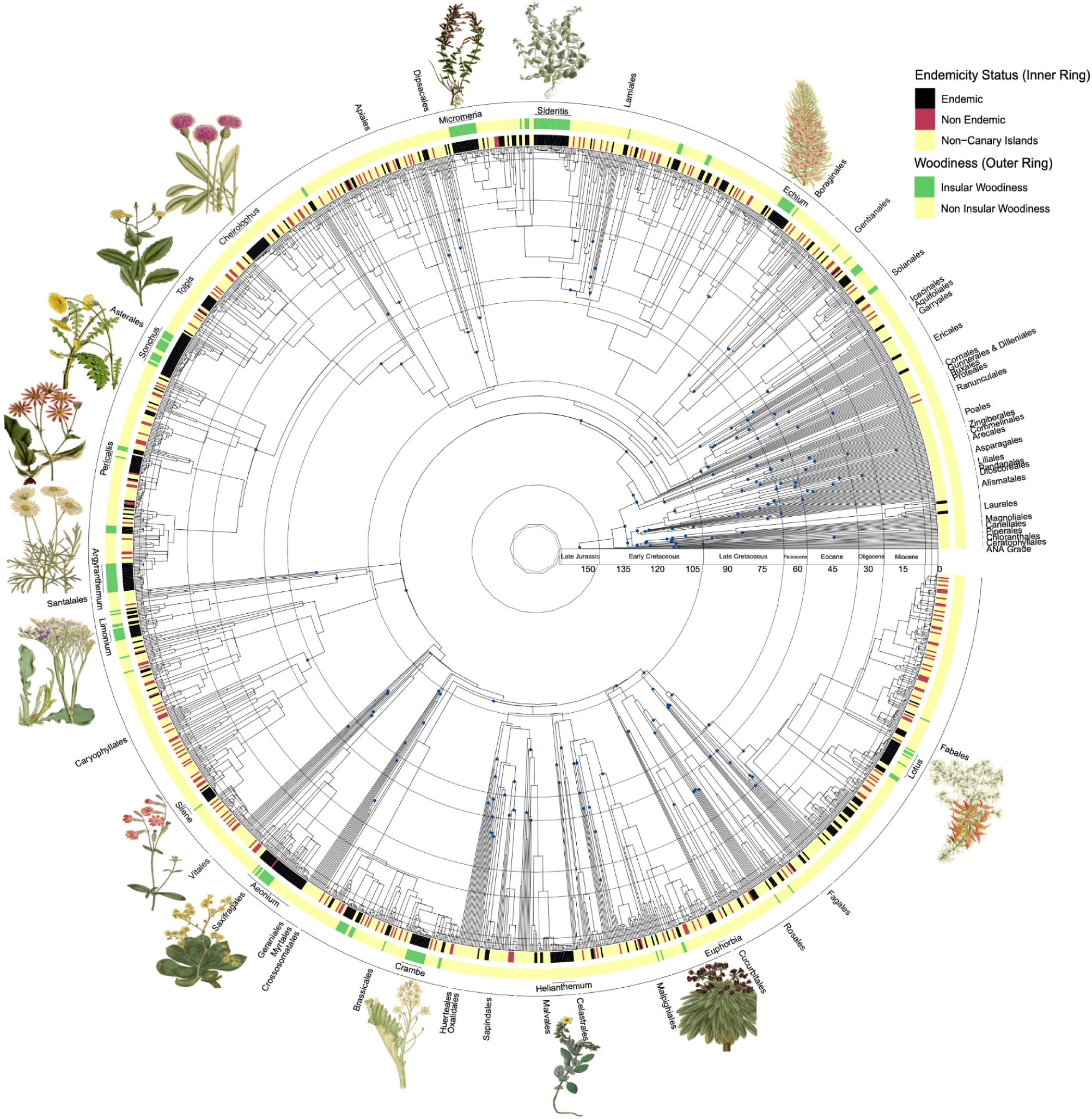
Time-calibrated phylogenetic tree for the Canary Islands non-monocot angiosperms, with the top 15 most species-rich lineages highlighted with illustrations. Taxonomic orders are given outside of the tree. The inner coloured ring denotes Canary Island endemicity status, with mainland relatives and additional taxa required for fossil calibration as ‘non-Canary Islands’. The outer ring shows the distribution of insular woodiness across the Canary Islands angiosperm tree. The blue circles at the nodes indicate fossil-calibrated nodes, given in Supplementary Data S7. The grey circles within the tree denote geological periods. For details on the illustrations, see Supplementary Data S10.

From our phylogenetic reconstruction, we recover many established relationships across the backbone of angiosperms and within Canarian lineages, alongside peculiarities (Figure 2, Supplementary Data S6). Major clades previously identified within angiosperms are recovered as monophyletic (ANA Grade, magnoliids-Chlorantales, monocots (and the magnoliid-monocot sister relationship recovered in (52)), and eudicots (super-asterids and -rosids)). The main peculiarity that affected the time estimation of a considerable number of Canarian lineages is the position of Caryophyllales (68 sequenced Canarian species, 51 lineages) and Santalales (1 sequenced Canarian species), which are retrieved as more closely related to rosids than asterids (cf. 52 but with Caryophyllales diverging earlier; see (60–62) for a position closer to asterids rather than rosids). In addition, as with (60–61), Vitales are recovered as sister to the rest of the rosids, rather than Saxifragales. As these peculiarities are towards the root of rosids/asterids, the topological differences can affect the colonisation and branching times obtained for Canarian lineages. However, changes in the backbone of our phylogeny do not affect the number of inferred colonisations.

We recover all large, charismatic radiations within the native non-monocot flora, comprising the most species-rich lineages, as monophyletic (Figure 2, Supplementary Data S6). Unexpectedly, the five endemic *Viola* species are largely recovered as independent lineages, with the only sister relationship retrieved being between *V. palmensis* and *V. odorata.* Interestingly, several native non-endemic Canary Island species are recovered as more closely related to Canarian congeners than to conspecifics from the continent. For example, *Clinopodium menthifolium* and *C. nepeta* specimens from the Canary Islands are recovered as more closely related to each other than to mainland populations of their own species (Supplementary Data S6). This pattern was also found in *Cuscuta, Rubia, Calendula, Filago, Senecio, Hippocrepis,* and *Ononis*. Whilst this pattern could be caused by multiple factors, such as incomplete lineage sorting or taxonomic gaps resulting from cryptic diversity, it is compatible with the surfing syngameon hypothesis (SSH: 45, 63), which proposes that intrageneric hybridisation between congeners becomes possible following recurrent island colonisation in phylogenetically closely related taxa previously geographically isolated on the mainland but occurring in sympatry on the island.

Insular woodiness is phylogenetically widespread in the Canary Islands, occurring in a total of 41 lineages from a diversity of orders and families. However, in terms of species diversity, IW is highly concentrated in a few lineages (Figure 2, outer ring). IW is present in 11 of the 15 most diverse island lineages, which account for 173 of the 241 IW species in the Canaries (72%; Supplementary Data S9, S11). Further, of the 41 lineages containing IW species, 24 have radiated into three or more species, pointing to a possible link between IW and diversification. For comparison, of the remaining 635 lineages without IW species, 31 have three or more species (Supplementary Data S1, S9). However, there are 11 IW singleton lineages demonstrating that IW is not always linked to speciation (Supplementary Data S11), with these singleton lineages not consistently recovered as the youngest IW lineages (i.e., younger IW lineages have diversified more; see Figure 3 below). The *Aeonium* Alliance (*Aeonium, Aichryson, Monanthes*, and former *Greenovia*; Crassulaceae) and *Sideritis* sect. *Marrubiastrum* (Lamiaceae) both have the most IW species in a single lineage, with 27 species each. These results align with those of (39), who found that although IW is widespread throughout non-monocot angiosperms, diversity is clustered within phylogenetically distantly related families.

**Figure 3.**
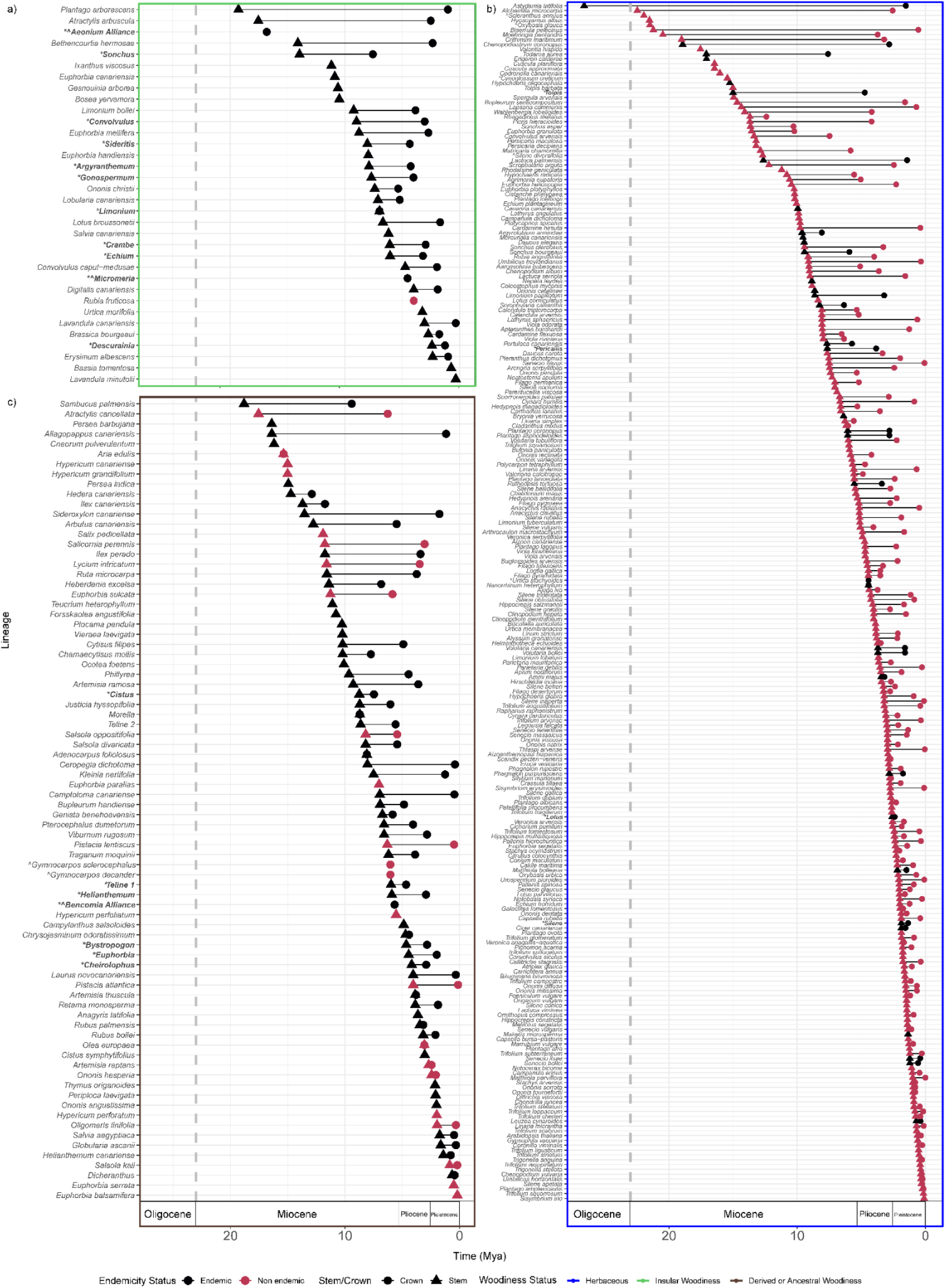
Colonisation intervals of Canary Island lineages, classified by growth form, with a) insular woody only (green), b) herbaceous (blue), and c) ancestrally and derived woody lineages combined (brown). Emboldened lineages with an asterisk (*) denote radiations (≥ 5 species). Lineages with a caret (^) indicate those for which their mainland relative was not successfully sequenced so the colonisation time may be overestimated; for singleton lineages, the stem age is given, for non-singleton lineages, the crown age is given. Overlapping stem and crown colonisation timed denote lineages recovered in our phylogenetic tree as polytomies. The vertical dashed line represents the age of the oldest extant Canary Islands rock (23 My). For lineages recovered as polyphyletic, a conservative estimate of age of the clade is used (see Supplementary Data S12).

### No evidence for clade-age hypothesis in Canary Island lineages

Based on our angiosperm-wide phylogeny that was time-calibrated with 166 fossils (Supplementary Data S7, see ‘Sampling Strategy’ in Methods), we estimate both stem (divergence from closest main-land relative on the tree) and crown ages (earliest branching event within the Canary Islands, applicable only to lineages that have undergone in-situ cladogenetic speciation). While there is much discussion about whether stem or crown age is the best proxy for colonisation time (64, 65), they provide an upper (stem) and lower (crown) bound for the colonisation event of a lineage. For cases where only a stem is available (lineages that have not diversified and are represented by a single tip in our tree), this can be assumed to be the maximum time of colonisation, where there is no lower bound. We estimated stem and crown ages for 362 Canary Island lineages (those from Figures 2, 3 for which we could confidently estimate colonisation time based on sequencing depth); for 291 of these lineages, there is no previously published colonisation estimate (Supplementary Data S12, S13). Across all lineages, the mean stem age is 6.6 million years (My) and the mean crown age is 4.4 My (Figure 3, Supplementary Data S12). Confidence intervals for each clade age estimate (crown and stem ages of Canary Island lineages) are given in Supplementary Data S14.

As we are particularly interested in the evolution of IW, we compared lineage ages between different growth forms. We define growth form as herbaceous, ancestrally and derived woody combined (i.e., lineages that colonised the islands as woody, hereafter ‘ancestrally+derived woody’), or insular woody (i.e., in-situ woodiness evolution on the islands; see (42) for further explanation of these growth forms). Woody island lineages (both ancestral+derived woody and insular woody) are generally recovered as older than herbaceous lineages (Figure 3). Mean stem and crown ages of insular woody lineages are 7.7 and 4.6 My, respectively (Figure 3a), whereas ancestrally+derived woody lineages combined have mean stem and crown ages of 7.9 and 5.4 My (Figure 3c). In comparison, mean stem and crown ages for herbaceous lineages are 5.7 and 3.5 My (Figure 3b). These comparatively younger mean ages of herbaceous insular lineages might reflect the increased ability of herbaceous species for long-distance dispersal required to colonise the Canaries (see Table 1), as well as the shift towards more open vegetation observed in the Pliocene Western Mediterranean (66, 67).

**Table 1.** Best fitting DAISIE models. Results are shown for the preferred background model (M4) and the two preferred differential rate two-type models (M26, M18). ‘λ^c^’-cladogenetic speciation; ‘µ’ - extinction rate; ‘*K*’ - carrying capacity; ‘γ’ - colonisation rate; ‘λ^a^’ - anagenetic speciation. V_2 of these parameters represents estimated rates for species of the ‘woodiness’ type in M26 and the ‘IW’ type in M18. ‘Prop’ is the proportion of the mainland pool of the “two-type” species (woody or IW). For explanations of how the proportions were calculated, see Methods and Supplementary Data S11. GF - growth form.

| Model Type | Model | $\lambda^c$ | $\mu$ | K | $\gamma$ | $\lambda^a$ | $\lambda^c\_2$ | $\mu\_2$ | K_2 | $\gamma\_2$ | $\lambda^a\_2$ | Prop | loglik | df | BIC |
| --- | --- | --- | --- | --- | --- | --- | --- | --- | --- | --- | --- | --- | --- | --- | --- |
| Background Model | M4 | 0.284 | 0.504 | Inf | 0.012 | 0 | - | - | - | - | - | - | -4904 | 3 | 9814.230 |
| Herbaceous vs Woodiness | M26 | 0.283 | 0.501 | Inf | 0.016 | 3.7E-07 | 0.283 | 0.347 | Inf | 0.001 | 3.70E-07 | 0.233 | -4749 | 6 | 9505.966 |
| Other GFs vs IW | M18 | 0.271 | 0.498 | Inf | 0.013 | 0 | 0.533 | 2.20E-13 | 51.212 | 0.001 | 6.63E-05 | 0.071 | -4851 | 8 | 9713.991 |

Whilst variance with existing estimates is expected given differences in sampling and dating methodologies used, our colonisation time intervals are consistently older than existing estimates. The mean stem and crown ages for the 73 Canarian angiosperm lineages with published colonisation time estimates are 5.5 My and 3.0 My, respectively (64, 65), while our mean colonisation times for these lineages are 7.7 and 4.2 My, respectively (Supplementary Data S13). Nevertheless, our consistently older divergence times do not radically change inferences on the biogeographic history of the Canary Islands. We find that all divergence times estimated here fall within the estimated age of the currently emerged islands of the archipelago (age of Fuerteventura, 23 My (68)), except *Astydamia latifolia* with an estimated stem age of 26.6 My. Additionally, our estimates of colonisation time intervals overlap with the late Miocene aridification and intensification of drought in the Western Mediterranean, Northern Africa, and Macaronesia, as previously reported (42, 64, 66, 69). These events, along with the onset of the Mediterranean climate and the Plio-Pleistocene glaciations, would have abruptly altered the Canarian climate and likely resulted in extinction events within the Canarian flora (42), driving taxon turnover. Taken together, these findings contribute to the growing body of evidence that extant Canarian angiosperm diversity likely reflects recent rapid taxon turnover mediated by high extinction and high colonisation rates (64).

To test the clade-age hypothesis, we investigated the link between the age of Canary Island lineages and species richness using a linear regression with the age estimates (both stem and crown) from our time-calibrated phylogenetic tree (Figure 4). In contrast to regional and global studies addressing the effect of clade age on species richness (6, 10, 18), we find no significant relationship between lineage age and species richness, except in the crown age dataset for IW lineages (albeit with low *R*^2^). These findings are also robust to grouping all growth forms (i.e., no significant relationship between lineage age and species richness; Figure S1), log-transforming the data, removing singleton lineages (both endemic and non-endemic), and accounting for phylogenetic non-independence using phylogenetic least squares regression. A lack of relationship between lineage age and richness can be evidence of spatio-ecological limits to diversity as well as high extinction rates within a system (10,64).

**Figure 4.**
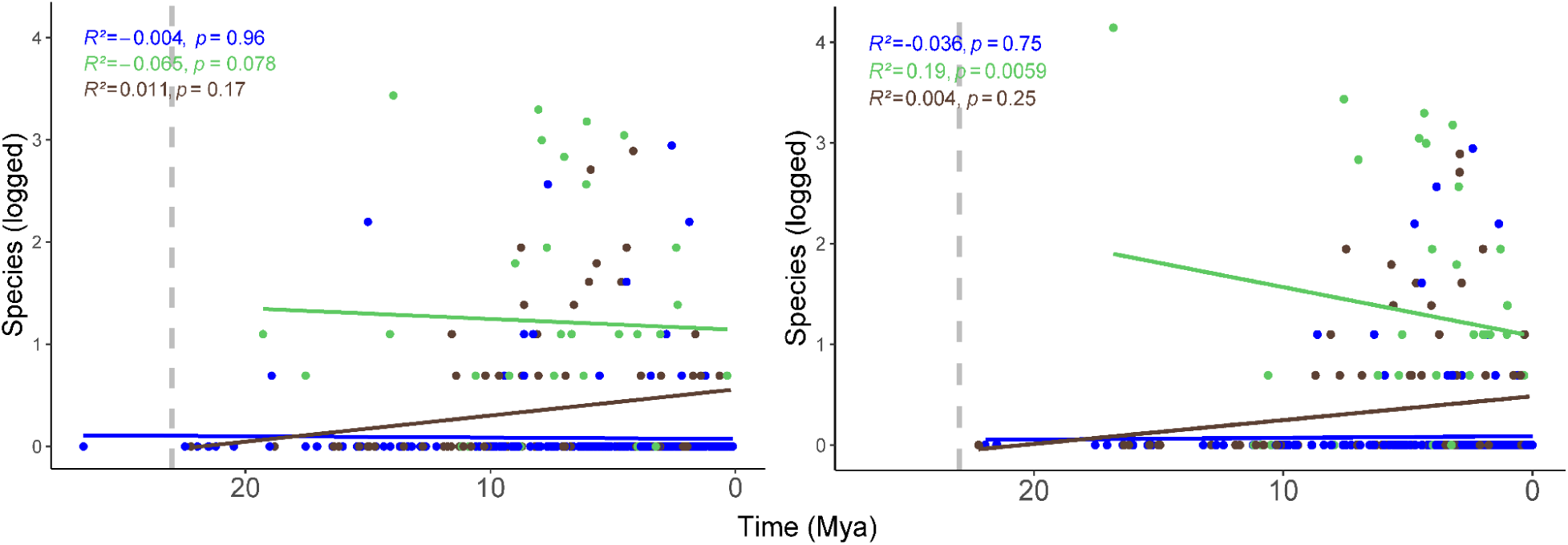
The relationship between lineage species richness and stem (left) and crown (right) age estimates. The correlations shown are for the unlogged richness estimates. The different colours refer to different growth forms: green is insular woodiness (n = 34), blue is herbaceousness (n = 248), and brown is ancestral and derived woodiness combined (n = 80; see Figure 3). The vertical dashed line represents the age of the oldest extant Canary Islands rock (23 My).

### Evidence for both diversification rate and spatio-ecological limits operating amongst lineages with different growth forms

To test the diversification rates and ecological limits hypotheses, we applied macroevolutionary dynamic stochastic modelling to our phylogenetic data. The colonisation and branching time data from our time-calibrated phylogeny were input into DAISIE, a phylogenetic model of island biogeography, to estimate biogeographical and diversification rates per lineage (i.e., speciation, extinction, and colonisation) for the flora of the Canary Islands. We fitted a set of 30 DAISIE models to our dataset (see Methods). Briefly, we fitted four models (M1-M4) that assume homogeneous rates of speciation, colonisation and (natural) extinction across the non-monocot angiosperm flora of the Canaries (background model in Table 1). In addition, to test whether growth form affects diversification rates, we fitted a set of “two-type” models (M5-M30), which allow for a subset of the non-monocot angiosperm flora to have evolutionary rates that differ from the background rates for the archipelago (14). Specifically, we tested whether lineages that were herbaceous vs woody at the time of colonisation have different diversification rates, and whether clades with IW species have different rates than clades without any IW species (see Methods for more details).

Table 1 shows the preferred DAISIE models for the background, the herbaceous vs woody, and the IW analyses. Under the preferred background model (M4), diversification in non-monocot angiosperms can be explained by just three parameters: cladogenetic speciation, extinction, and colonisation. Under the M4 model, the system is diversity-independent, that is, we find no evidence that diversity leads to a slowdown in rates of colonisation or speciation, and no detectable upper bound to clade-level diversity on the Canaries. Furthermore, the model indicates that the system acts as an evolutionary sink - the archipelago needs to receive new colonists from the mainland to maintain diversity, given that the rate of extinction greatly exceeds speciation. The colonisation rate is exceptionally high compared to other studies, at approximately 234 colonisations per million years (My; gamma * mainland pool size; 0.012 * 19,631), further supporting the findings of Garcia-Verdugo et al. (64) that extant Canarian flora is maintained by high extinction and high colonisation rates, as predicted by the SSH (45, 63). In comparison, Canary Islands land birds have been estimated to have 45 colonisations per My (70). Under the background model, the diversity of Canary Islands non-monocot angiosperms is found to be at a dynamic diversity equilibrium (diversity remains relatively constant through time, Figure S2), as was the case in Canary Islands birds (70), in contrast with angiosperm diversity globally (71). Finally, whilst the presence of many lineages with just a single endemic species would suggest that rates of anagenetic speciation (without lineage splitting within the archipelago) are high, the DAISIE analysis favours a scenario where such lineages are, surprisingly, exclusively the result of cladogenetic speciation followed by extinction.

We found that models that allow differential rates based on growth form are preferred over the background models. For the herbaceous vs woodiness analysis, the preferred model was M26 (Table 1). Under this model, woody species at the time of colonisation (including ancestrally woody and derived woody, but not IW species) have a lower extinction rate than species that were herbaceous at the time of colonisation. However, they have the same speciation rates, contrasting with previous angiosperm-wide diversification studies that found that herbaceous species with shorter generation times have higher rates of molecular evolution and therefore higher speciation rates (72, 73). Furthermore, under this model, differences in diversity between herbaceous and woody species are explained by pronounced differences in colonisation rates. Where the rate of colonisation for herbaceous species is estimated to be 0.016, equivalent to 307.6 colonisations per My (0.016 * 19,631), the rate of colonisation for woody species is only 0.0001, equivalent to 0.6 colonisations per My (gamma * mainland pool size * type 2 proportion; 0.00012 * 19,631 * 0.23). Improved colonisation capabilities of herbaceous compared to woody colonisers on oceanic islands have been posited since the time of Darwin (20), largely due to expected differences in propagule size (74). Generally, small-seeded species are better dispersers than large-seeded species once plant height is accounted for (75, 76), but a differential ability of herbaceous species to colonise islands, relative to woody species, based solely on growth form, remains hitherto untested.

For the analyses of IW versus other growth forms, we find that parameters are different for lineages with the potential to evolve IW under the preferred model (M18; Table 1). Both cladogenetic and anagenetic speciation are higher for IW lineages, whereas extinction is substantially lower. The colonisation rate of IW lineages is also substantially lower than the colonisation rate for non-IW lineages, at 0.18 colonisation events per My (gamma * mainland pool size * type 2 proportion; 0.00013 * 19,631 * 0.0706), meaning the diversity of IW lineages on the Canaries is driven by high speciation on the island, rather than the sink dynamics of the background community. This result supports the diversification rate hypothesis to explain differences in diversity between IW lineages and the background community. Furthermore, IW lineages are identified as having diversity-dependence in rates of colonisation and cladogenesis, in support of the ecological limits to diversity hypothesis. Under this scenario, the more IW species there are on the island, the lower the rates of colonisation and cladogenesis for IW lineages. This is not the case for non-IW species, for which no diversity-dependence is detected. The causes for the evolution of IW are incompletely understood, with drought resistance, favourable climate and reduced herbivory (selecting for longevity), competition for light (thereby selecting for increasing height), and burial avoidance following volcanism, are all suggested global drivers that may interact with each other in a different way for each lineage, often acting lineage-specifically (39, 40, 77). Under the preferred models for both growth form analyses (M26, M18), the inference of equilibrium in Canarian non-monocot angiosperms was unchanged (Figure S2). Finally, though we favour using BIC to identify model preference among DAISIE models due to lower error rates compared to AIC (15), when using AIC, the inference of differential rates for IW lineages is unchanged under the preferred models. Using AIC, M26 is identified as the preferred model, as with herbaceous vs woodiness analysis, with IW lineages having a dramatically reduced extinction and colonisation rates relative to non-IW lineages.

In conclusion, using our phylogenomic data, we have shown that of the three main global hypotheses to explain global biodiversity patterns, there is strong evidence for the diversification rate hypothesis and evidence for the spatio-ecological limits hypothesis. First, we find that the flora of the Canary Islands is shaped by high extinction and colonisation rates (Table 1) and that there is no relationship between lineage age and diversity, supporting a scenario with a high turnover of lineages (63, 64), and rejecting the clade age hypothesis (Figure 4). Second, IW lineages behave uniquely compared to the total archipelago-wide flora of the Canary Islands, with higher speciation and lower extinction rates than the rest of the non-monocot flora, supporting the diversification rate hypothesis to explain unevenness in diversity. This hypothesis is also supported by our finding that woody species have reduced extinction rates relative to herbaceous species. (Table 1). Third, IW lineages have a detectable carrying capacity, meaning that as the number of IW species increases, rates of cladogenesis and colonisation decline (Table 1) and we show that the Canarian non-monocot angiosperm flora has achieved a macroevolutionary equilibrium in terms of species richness (Table 1, Figure S2), supporting the spatio-ecological limits hypothesis. With our time-calibrated phylogeny, many new avenues of research into the Canary Islands angiosperm flora become possible, including on trait(-dependent) evolution (78), island-wide vs clade-specific diversification (79), and evolutionary return time (34, 80), whilst the archipelago checklist identifies the remaining lineages to be sequenced in order to complete the tree of life for this iconic flora (Supplementary Data S1).

## Methodology

### Canary Islands as a Model System

The Canary Islands are an archipelago comprised of seven main islands (El Hierro, Fuerteventura, Gran Canaria, La Gomera, La Palma, Lanzarote, and Tenerife), two smaller islands under 30 km^2^ (La Graciosa and Alegranza), and several smaller islets. The oldest extant exposed rock emerged ca. 23 million years (My) ago (68), with many more volcanic islands previously present over the last 70 My, which have since become submerged (53, 74, 81). Located in the Macaronesian meta-archipelago off the northwest coast of Africa, at the easternmost point, the Canary Islands lie only approximately 100 km off the coast of Western Sahara.

Alongside Hawai’i and the Galápagos Islands, the Canary Islands form one of the best-studied oceanic island systems (82), and the native angiosperm flora is among the best-studied island floras globally (45, 47). With an estimated ∼1,300 native species of angiosperms (53, 54, 74), the flora is the most species-rich of any oceanic archipelago (48), contains many evolutionary radiations (27), and holds the highest number of insular woody species and shifts towards insular woodiness globally (39, 41, 42).

### Checklist of native Canary Island angiosperm species

We compiled a list of all known native Canary Island angiosperms based on the most up-to-date published checklists (53, 54). From these we compiled all species in the following categories: Surely Native, Possibly Native (53); Nativo Seguro [Native with Certainty], Nativo Probable [Probably Native], Nativo Posible [Possibly Native] (54). We further refined the resulting list using expert knowledge to remove species that were deemed to have been erroneously considered native (53-55, 83; Supplementary Data S1). For each species, we recorded their endemicity status (endemic (occurring only in the Canary Islands) or non-endemic (occurring natively in the Canary Islands and elsewhere)) and their growth form (annual herbaceous, perennial herbaceous, insular woodiness, derived woodiness, or ancestral woodiness) using published literature and floras (55, 84–89). Biennial herbaceous species were reclassified to ‘perennial herbaceous’ to give an annual and non-annual herbaceous category. For the specific reference used for each species, see Supplementary Data S9.

### Estimating angiosperm colonisations of the Canary Islands

Using our updated checklist of Canary Island angiosperms, we estimated the number of times each natively occurring genus colonised the Canary Islands and gave a written description for each inferred colonisation event. This step was essential for designing the sampling strategy for our phylogenetic reconstruction (see ‘Sampling Strategy for our Phylogenetic Tree’ below), but was also used to compute metrics for lineages that were not included in the phylogeny (see ‘Statistical Analysis of Colonisation Data’ below). Whenever prior phylogenetic information was available, we used phylogenetic trees to infer the closest mainland relative of each native Canary Island angiosperm lineage. For a list of the phylogenies used for each genus, see Supplementary Data S1, and for a list of the mainland relatives used per Canary Islands lineage, see Supplementary Data S2. A genus was considered to have colonised the Canary Islands several times if it had multiple species native to the Canaries which do not cluster together in the phylogeny. For example, if two Canarian species are more closely related to different mainland species than each other, they were counted as two colonisations. It is important to note that our method does not account for multiple colonisations of the Canary Islands by the same species. While these are known to occur (45,63,90), this information is not available for most species and can be difficult to infer from molecular data. Therefore, we chose to focus on colonisations at the genus level. Our final estimate is, therefore, an underestimate of the total number of angiosperm colonisations on the Canary Islands, but a good estimate of the number of colonisations that gave rise to extant Canary Island lineages.

For genera for which no well-sampled phylogenies were available (e.g. did not include all Canarian species, or did not sample mainland species presumed to be close relatives), we applied a measure of uncertainty for the number of colonisations. A minimum bound counted how many distinct colonisations were possible to estimate using the available literature and a maximum bound assumed that every species colonised the Canary Islands separately. A ‘likely’ number of colonisations for each genus was also estimated using available phylogenetic literature, distribution data, and the endemicity status of the native Canary Island species. For example, if a genus had multiple native Canarian species, with several endemics and several island species that are also widespread on the continent, the endemic species were assumed to result from a single colonisation in the ‘likely’ estimate and the non-endemics were considered to result from a single colonisation each. Our final estimate of Canary Island angiosperm colonisations thus consisted of a minimum, likely, and maximum number of colonisations. For full explanations for each inferred colonisation event, see Supplementary Data S1, ‘Inference Notes’.

### Statistical Analysis of Colonisation Data

From the previous steps, we obtained the total number of Canary Island species and likely colonisations for each family of flowering plants represented in the Canary Island flora. We assessed the relationships between the native mainland species richness, the number of Canary Island colonisations for each family, and the number of native Canary Island species. We downloaded native angiosperm species diversity data for the countries comprising the Mediterranean Basin and Macaronesian enclave (including the Mediterranean islands) from the World Checklist of Vascular Plants using Version 12 (28 September 2023) from the R package ‘rWCVPdata’ (91). Though not all native species and families on the Canaries have colonised from the Mediterranean Basin, 74 of 83 families occurring on the Canary Islands are present in the countries that comprise the Mediterranean Basin. These 74 families comprise 1,358 (possibly) native species on the Canaries (Supplementary Data S4), meaning we capture a nearly complete representation of Canary Island species using the Mediterranean Basin as the comparative mainland. Additionally, the Canarian flora has a close affinity with the Mediterranean flora (92), and as the limits of the Mediterranean Basin fluctuate over time, this explains the close affinity of these floras, justifying the use of the Mediterranean Basin as our choice of mainland system.

We assessed correlations between Canary Islands angiosperm richness, number of colonisations, and native mainland species richness at the family level. All data were log-transformed before computing Kendall’s rank correlation coefficient. As the subsequent dataset contained many taxonomic groups with < 5 species present on either the Canary Islands or the mainland, these taxa were removed to avoid biasing the data. All reported values are subsequently given for the filtered dataset containing ≥ 5 species, although these relationships were identical to those using the full dataset. Island disharmony in comparison to the Mediterranean mainland was assessed using binomial tests to determine whether family-level native angiosperm richness on the Canary Islands was significantly under-/over-represented relative to the Mediterranean Basin (35). Binomial tests were run for each family using the ‘binom_test()’ function in R, where *x* represents the total number of native angiosperm species, *n* represents the number of native mainland species, and *p* represents the proportion of island species relative to the total number of angiosperms native to the Mediterranean Basin.

### Sampling Strategy for Our Phylogenetic Tree

In this section, we describe the sampling of plant specimens used for our time-calibrated phylogeny. A fully sampled tree would include all 1,385 species of Canary Island angiosperms as well as representatives of each of their closest extant mainland relatives, which would allow estimating stem and crown ages for all Canary Island non-monocot lineages. However, such a high level of sampling is currently not feasible using our approach because, for many Canary Island lineages, it is currently not possible to infer their closest mainland relative (e.g., Canary Island species belonging to genera that are very diverse on the continent, and for which no well-sampled published phylogeny is available), or because good quality samples for DNA sequencing are not available in herbaria. Based on our updated phylogenetically informed checklist of Canary Island angiosperms, we were able to infer the number of colonisations and identify the closest mainland relatives for the majority of Canary Island lineages. We used this information to guide our sampling for the new phylogenomic dataset produced for this study.

We chose to focus on non-monocots because monocots never produce a wood cylinder due to the absence of a vascular cambium (41, 51); as we are interested in the effects of IW on diversification, monocot lineages were removed from the sample so as not to bias our analyses against the presence of woodiness. Our sampling group for the new phylogeny thus consisted of every native, non-monocot Canary Islands angiosperm species for which we could identify the nearest mainland ancestor and the number of colonisations could be estimated (see Supplementary Data S1). This consisted of 838 native Canarian angiosperm species (∼61% of the total native Canary Islands angiosperm species) comprising 437 colonisation events (50%), and two outgroup species (the most closely related extant mainland taxon) for every native Canarian species, totalling 1,710 specimens to sample for this study (Supplementary Data S2). A recent simulation study has shown that this sampling fraction and our approach, by including the majority of species-rich lineages (24/30 (80%) of the most species-rich angiosperm genera, or 25/30 (83%) of the most species-rich non-monocot lineages), leads to very little loss of information compared to complete sampling using DAISIE to estimate biogeographical parameters (93, In press). Our sampling was consistent with existing studies interested in obtaining colonisation estimates (65), consisting of one individual per native Canary Island species and two outgroup species per lineage. For the outgroup species, we sampled the two most closely related mainland species for Canary Islands endemics, and for non-endemic, native species we sampled two different nearby mainland populations.

To generate our phylogeny, we used a combination of existing Angiosperms353 sequence data and newly generated sequences from fresh, herbarium, and DNA-bank material using capture-based hybridisation sequencing. For this study, we obtained dried leaf material specimens from 23 herbaria and botanical gardens throughout Eurasia and the Canary Islands, as well as from colleagues collecting field samples (Supplementary Data S12). We generated new sequences for 1,244 specimens (495 native Canarian species and 749 outgroup taxa; Supplementary Data S2) of the 1,710 target specimens, representing 735 unique species. Previously published sequence data were also obtained for 210 specimens: 25 *Helianthemum* (58), 22 Brassicaceae (57, unpublished data), and 163 Asteraceae (59) from co-authors. To maximise the number of nodes that could be calibrated using fossils in our analysis, we complemented our dataset with publicly available data from the Plant and Fungal Tree of Life Project (PAFTOL; 291 samples; 49,52), for a total of 1,745 tips. Seven gymnosperm species were used to form the outgroup, one from each recognised order, except Welwitschiales. See Supplementary Data S15 for all specimens included in the study.

### DNA Extraction, Library Preparation, Target Capture, and Sequencing

For all herbarium or field collected samples, we extracted genomic DNA from an average of 20 mg of leaf material (or branch, stem, and/or flowers if no leaf material was available or if the species did not produce leaves) using the DNeasy PowerPlant Pro Kit (Qiagen, Hilden, Germany), following manufacturers protocols but repeating the final elution in two x 60 ul elutions rather than one 120 ul elution. DNA concentrations were quantified using the Agilent 5300 Fragment Analyzer with the High Sensitivity Genomic DNA Analysis Kit (DNF-488; Agilent Technologies, Santa Clara, CA, USA). Genomic DNA was stored at the Naturalis Biodiversity Center, Leiden, The Netherlands.

DNA extracts with an average fragment length > 400 bp were sheared using sonication in an M220 Focused-ultrasonicator (Covaris, Woburn, Massachusetts, USA) to obtain an average fragment length of 350 bp. Genomic libraries were prepared using the NEBNext Ultra II DNA Library Prep Kit (New England Biolabs, Ipswich, Massachusetts, USA). This process consisted of end repair of sheared or fragmented DNA, adaptor ligation, size selection, and PCR enrichment of adaptor-ligated DNA over 8-14 cycles. The quality of the libraries was then assessed using the Agilent 5300 Fragment Analyzer HS Small Fragment DNF-477 kit. Indexing was conducted using 384 unique combinations of IDT10 primers (Integrated DNA Technologies, Coralville, Iowa, USA), following the methodology of Hendriks (94).

The libraries were pooled into groups of 7-40 specimens based on total DNA quantity (ng), fragment length distribution, and taxonomy (to the order level). Pools were enriched using the Angiosperms353 bait set (Arbor Biosciences myBaits® Target Sequence Capture Kit, Arbor Biosciences, Ann Arbor, Michigan, USA; hereafter ‘A353’; 95), targeting 353 genes, following the manufacturer’s recommendations. An experimental pool of 62 specimens was also enriched using the A353 bait set to test the efficacy of enriching large pools of specimens distantly related by taxonomy. Hybridisation was performed at 65°C for 24 hours for the first 176 specimens, but was reduced to 62°C for the remaining specimens, as we found that this increased the percentage of loci on target recovered. Post-hybridisation libraries were amplified using KAPA HiFi HotStart Ready Mix (Roche Sequencing) for 13-32 cycles, followed by a bead clean-up. Amplified libraries were then ‘spiked’ with 0.2 ul of unenriched library pool (pre-capture pools). To reduce sequencing costs, amplified libraries were then pooled equimolarly into 19 endpools, avoiding duplicated index combinations. Sequencing was performed using a NovaSeq 6000 sequencer (Illumina, San Diego, California, USA) at BaseClear, the Netherlands, producing 150 bp paired-end reads at 100x targeted coverage.

### Sequence Assembly and Paralog Filtering

Raw, demultiplexed sequence data, in the form of FASTQ files, were quality-controlled and trimmed using Trimmomatic v0.39 (96), using the settings of Baker et al. (49). Trimmed reads were then mapped using HybPiper version 2.1.6 using the ‘mega353’ target file (97, 98). The BWA algorithm was preferentially used over BLASTX as this resulted in more reads mapping for our samples (99,100 but see Murphy et al. (101)). Subsequently, each gene was assembled using SPAdes version 3.15.3 and coding sequences were extracted using Exonerate version 2.4.0 (102, 103).

Three summary functions provided by HybPiper were run: ‘hybpiper stats’, ‘retrieve_sequences’, and ‘retrieve_paralogs’. HybPiper stats provides summary statistics related to mapping success, including the percentage of reads on target, the number of genes with sequences, and the number of genes with sequences of 25-150% of the target length. This was run for all samples to gauge the success of mapping and contributed to whether a specimen needed to be resampled (see Supplementary Data S15 for HybPiper stats summary of the final specimens used in this study). Retrieve_sequences pulls the best (unfiltered) contig sequence fasta files (in .FNA format) for each locus, regardless of the paralogs warning. Similarly, retrieve_paralogs pulls the mapped sequence fasta files flagged with a paralog warning; if no paralog warning was detected, the best contig is retrieved from the HybPiper output. Phylogenetic analyses including paralogous genes (homologous genes that share a common ancestor as a product of gene duplication) can produce incorrect topologies (104, 105). Given the prevalence of paralogs in many of the families included in our dataset (i.e., Brassicaceae and Asteraceae; 57, 106), and the presence of several groups which are known to hybridise in our dataset (thereby increasing the likelihood of paralogs being present via allopolyploidy; 104, 107, 108), paralogs were filtered to assess the effect on tree resolution and branch support.

Once the mapped contigs and paralogs had been retrieved, we used ParaGone to filter putative paralogs (104, 109). To identify the best-fitting ParaGone algorithm for our dataset, each algorithm was run (Maximum Inclusion (hereafter ‘MI’), Rooted Ingroups (‘RT’), and Monophyletic Outgroups (‘MO’); see Yang and Smith (109) for further details), and phylogenetic reconstruction was performed using the orthologs output from each algorithm (see below).

### Gene Alignments, Tree Reconstructions, and Time Calibration

Once the orthologs had been retrieved from ParaGone, gene alignments were generated separately for each gene using OMM_MACSE (110, 111). OMM_MACSE was used over MACSE for its ability to better handle large numbers of sequences whilst still generating codon-aware alignments, by utilising external alignment software that does not produce codon-aware alignments, such as MAFFT. Aligned genes were then trimmed using TrimAL v1.3 (112), with the parameters resoverlap 0.75, seqoverlap 0.64, and gt 0.70. Gene trees were generated using maximum likelihood methods from the trimmed alignments using ultrafast bootstrapping in IQ-TREE v.2.2.2.7 (113, 114). These steps were repeated, using the same parameters, for the unfiltered contigs obtained from the retrieve_sequences function.

We used three approaches to compare the effect of paralog filtering on phylogenomic tree reconstruction. First, we used the gene trees from IQ-TREE generated from the unfiltered contigs to build multi-species coalescent species trees using ASTRAL-IV v1.18.4.5 and ASTRAL-Pro3 v1.18.3.5 (115, 116). A preliminary species tree was produced using ASTRAL-IV to identify specimens with obviously erroneous placement (i.e., recovered completely incorrectly due to specimen misidentification, contamination, or very poor-quality input data); these specimens were then resampled (390 specimens). The wet lab, sequence assembly, and gene alignment steps were repeated for resampled specimens and a new non-paralog filtered multi-species coalescent species tree was produced via ASTRAL-IV to re-assess erroneous placement. Any specimens identified as erroneous after resampling were removed from our dataset to avoid incorrect inference in downstream analyses and the ASTRAL-IV tree was rebuilt (153 specimens). This process was conducted for both the non-paralog-filtered and paralog-filtered datasets.

Second, orthologous gene trees produced by each ParaGone algorithm were used to reconstruct multispecies coalescent species trees in both ASTRAL-IV and ASTRAL-Pro3. As each ParaGone algorithm produces a different number of orthologous groups, and resultantly, produces different topologies and node support values (117), we assessed the relative performance of each approach and compared the produced trees with those of the unfiltered ASTRAL-IV and ASTRAL-Pro3 trees. As the MI and RT algorithms produced too many orthologous groups to feasibly generate a species tree from (at 22,184 and 5,612, respectively), the MO algorithm was used to produce the paralog-filtered tree (which produced 111 orthologous groups). For our dataset, as paralogy resolution resulted in many more polytomies (see Supplementary Data S16), we proceeded with the non-paralog filtered ASTRAL-IV tree for the time calibration.

Third, we used a maximum likelihood (ML) supermatrix approach in RAxML-NG v1.2.1 to construct a phylogeny tree with branch lengths that could be used for divergence-time estimation (118). We adopted a ‘gene shopping’ approach, implemented in SortaDate (119), where gene trees were filtered first based on similarity to the species tree (the non-paralog filtered ASTRAL-IV tree), second by clock-likeness (root-to-tip variance), and then by tree length. Gene tree concordance was favoured as high gene tree conflict can lead to errors in divergence estimates (120). The best 25 loci were identified and used for the subsequent dating analysis. The trimmed alignments for these genes were concatenated using CatSequences (https://github.com/ChrisCreevey/catsequences) and RAxML-NG was run using the concatenated supermatrix with a GTR+G model, with 1,000 bootstraps, constraining the topology to the non-paralog filtered ASTRAL-IV tree. Divergence times were estimated using the relative rate framework implemented in RelTime for its increased accuracy relative to TreePL (121, 122), especially in the presence of convergent substitution rate shifts among lineages, which we expect within our dataset. Fossil calibrations were based on 163 unique minimum age calibrations from the AngioCal v.1.1 dataset, two Asteraceae-specific minimum age calibrations, and an updated age for crown Eudicots (50, 52, 123, 124; Supplementary Data S7). A maximum constraint of 154 million years was used at the crown node of angiosperms, as the resulting age estimates were found to match existing divergence estimates of many angiosperm families (52).

### DAISIE Analysis

We use DAISIE to estimate evolutionary and biogeographical rates of Canarian non-monocot angiosperms (14). DAISIE utilises phylogenetic information to estimate per lineage rates of colonisation (of an island system), cladogenetic speciation (speciation with lineage splitting within the island, e.g. leading to radiations), anagenetic speciation (evolution of new endemic species with no lineage splitting, does not lead to increased diversity on the island), and extinction, as well as diversity limits of a system (14, 15).

A DAISIE analysis is centred on a focal island assemblage, in our case all species and lineages of non-monocot angiosperms that occur natively in the Canary Islands. For each lineage, DAISIE requires its endemicity status, number of species, as well as its colonisation time and branching times (if any, in case of radiations). Note that the entire focal group needs to be considered, so both sampled and unsampled taxa need to be included. For all lineages sampled in our phylogeny, we extracted their information from our species checklist (endemicity, number of species) and the time-calibrated phylogeny (divergence times). Stem ages for these lineages were manually extracted from the time-calibrated ML tree, along with all branching times within Canary Island lineages. We treated these stem ages as precise ages of colonisation in the DAISIE analyses for endemic lineages, though these stem ages may better reflect divergence times than colonisation times in lineages with high extinction rates. For non-endemic lineages, these stem ages are treated as maximum ages, as we cannot be certain which mainland population colonised the Canary Islands; maximum ages reflect this uncertainty in colonisation estimate. As crown estimates have also been proposed as an adequate proxy for island colonisation time for endemic island clades (64, 65), we built another dataset with crown ages for Canary Island endemic species of the same monophyletic group, and stem ages for all other lineages (stem+crown dataset). In this dataset, endemic lineages were treated as precise ages of colonisation and non-endemic lineages as maximum ages (Supplementary Data S17). To avoid biasing against insular diversification, Macaronesian endemics were treated as Canarian endemics in the analyses (see Supplementary Data S1 for Macaronesian endemics), under the assumption that diversification happened within the Macaronesian meta-archipelago. Endemicity status in Figures 2 and 3 reflects this. Species that belong to our focal group but which are not sampled in the phylogeny (were not targeted for sequencing, or sequencing was not successful) are also included in the DAISIE analyses using the following approaches: a) if a species is not sampled in the phylogeny but is known to belong to a lineage that is sampled in the phylogeny, we add it as missing species to that clade, using the “missing species” option in DAISIE; b) if an entire lineage is missing from the phylogeny (can be a singleton lineage or a lineage composed of multiple species), we add this to DAISIE by specifying that the lineage could have colonised any time between the island age and the present; c) for lineages for which only the nearest mainland relatives were successfully sequenced, we used the stem age of the mainland relatives as a maximum age of colonisation. For three genera that were recovered as polyphyletic (*Cuscuta, Persicaria,* and *Viola*), the stem age of the oldest genus lineage was used as the maximum age.

DAISIE also requires an estimate of the size of the mainland pool of species that could have colonised the Canary Islands, to infer colonisation rates, and of the age of the insular system. The mainland pool size was estimated using native non-monocot angiosperm species data for the Mediterranean Basin from the World Checklist of Vascular Plants (WCVP) using the R package ‘rWCVPdata’ (91). A species list was built using native angiosperm species from all countries in the Mediterranean Basin. Monocot species, all duplicate instances per species, and introduced species were removed to give the number of native non-monocot angiosperm species in the Mediterranean Basin. Inaccuracies in the mainland pool size will affect inferred colonisation rates, but will not affect inferred speciation and extinction rates and subsequent model choice. The maximum age of the Canary Islands was fixed to 23 My following Campeny et al. (68).

We fitted two sets of DAISIE models to each stem and stem+crown datasets, using the DAISIE_ML function in the DAISIE R package (14, 125): background and two-type models. The background models estimate the overall background parameters that apply to the entire focal assemblage; two-type models allow a specified group of species of a certain ‘type’ to be governed by different values of parameters compared to the background. For the background models, we fitted a set of four models that varied in whether *K* (carrying capacity) was estimated (diversity-dependent model) or fixed to infinity (diversity-independent model), and in whether anagenesis was estimated or fixed to zero. For the two-type models, we assessed model fit in two separate growth form analyses: herbaceousness at the time of colonisation (i.e., herbaceous and insular woody species on the Canaries) vs woodiness at the time of colonisation (ancestrally woody and derived woody species; Supplementary Data 12), and insular woodiness vs all other growth form types (herbaceous, ancestrally woody, and derived woody) on the Canaries. For each of these growth form comparisons, we used the models 5-30 of Michielsen et al. (34), which vary one or more parameters per model (i.e., cladogenesis, extinction, carrying capacity (*K*), colonisation, and/or anagenesis). For a full description of each of the 30 models, see Michielsen et al. (34). Therefore, for each dataset (stem and stem+crown), we fitted a total of 30 models. We ran each of these 30 models 33 times, randomising input starting parameters. We then calculated the Bayesian Information Criterion (BIC) and Akaike Information Criterion (AIC) for each optimisation to select a preferred model. We favoured the BIC results as BIC has been shown to produce lower error rates when choosing among DAISIE models (15).

For the two-type analyses in DAISIE, we specified a proportion of species on the mainland that belong to each type. For the herbaceous vs woody analysis, the proportion of the type-2 species was calculated by combining three global datasets on growth form (85, 91, 126). The combined dataset was then filtered to only include the species in our mainland pool, and the growth form was retained for each dataset. As the WCVP does not record woodiness status, growth forms were reclassified to give binary woodiness/non-woodiness. To reconcile differences in the woodiness status of species between the three datasets, the average number of herbaceous and woody species was used to find the proportion for the herbaceous vs woody analysis (11,749 to 3,569, respectively). For the analyses of IW vs other growth forms, the mainland proportion was estimated by summing the number of herbaceous Mediterranean species in the most recent common ancestral lineage of the Canarian IW lineages (including those which are not majority IW but have IW species; see Supplementary Data S11 for specific lineage used). This method estimated 1,386 potentially IW species, or 7.1% of the mainland pool size (see Supplementary Data S11). To assess how this estimate could affect the results, we reran the analysis by incrementally varying the proportion by half to double our obtained estimate and the model choice remained consistent (results not shown). As model preference did not differ between the stem or stem+crown datasets, we report results only for the stem dataset, which is more comparable with previous studies using DAISIE, which mostly use the stem as the node for colonisation time proxy. Once the best model was identified for each growth form analysis, we used the DAISIE_sim function to simulate 1000 islands for the stem and stem+crown background and two type models.

## Supporting information

Supplementary Files Description

## Acknowledgements

We thank Lizzie Roeble (Naturalis Biodiversity Center) and Lucas Miller (Technical University of Munich) for their contributions to target group identification, and the following institutes/people who helped us with the sampling: Universidad de La Laguna (TFC); Jardín Botánico Canario Viera y Clavijo-Unidad Asociada de I+D+i al CSIC (JBCVC-UACSIC); Banco de ADN de la Flora Canaria of the JBCVC-UACSIC; Flora Iberica (CGL2005-05471-C04-01); Javier Morente-López (Goethe-University Frankfurt); Universidad de Sevilla (SEV); Steven Janssens (Meise Botanic Gardens, BR); Georgian Academy of Sciences; Royal Botanic Gardens, Kew (K); Museum National d’Histoire Naturelle (P), P gives access to the collections in the framework of the RECOLNET national Research Infrastructure; Suzanne Cubey (Royal Botanical Gardens Edinburgh E); Ranee Prakash (Natural History Museum, London); Roderick Bouman (Hortus Botanicus, Leiden); Leopoldo Medina (Real Jardín Botánico de Madrid, MA); Institute of Biotechnology of the National Academy of Sciences of the Krygyz Republic; Andreas Berger (Naturhistorisches Museum Wien, W); Henry Väre (The Botanical Museum of the Finnish Museum of Natural History, H); Patrik Frödén and Arne Thell (Lund University Botanical Museum, LD); Filipe Covelo (Herbarium of the University of Coimbra, COI); Reto Nyffeler (United Herbaria, Z & ZT); and Roxali Bijmoer, Susana Arias Guerrero and Marnel Scherrenberg (Naturalis Biodiversity Center). We also thank our Naturalis colleagues Kasper Hendriks, Sander van Zon, Marina Ventayol Garcia and Elza Duijm for their assistance with sequencing and wet lab work, and Arjan Schieven, Kasper Hendriks, Lizzie Roeble, Nina Kerdiles, and Edwin van Haas for their assistance with bioinformatics. We thank Abelardo Aparicio, Rafael González Albaladejo, and Juan Viruel for sharing the *Helianthemum* sequencing data (BioProject PRJNA1219060). The collection and sequencing of *Helianthemum* were funded through the following projects: PID2020-116355GB-I00 and CGL2017-82465-P. Most of the Asteraceae family was sampled and sequenced via the support of the following Spanish Ministry of Science and Innovation (MICINN)’s projects: ASTERALIEN - PID2019-110538GA-I00 and DecodAdapt - PID2023-147122NB-I00. We appreciate help from the Center for Information Technology of the University of Groningen to access the Hábrók high-performance computing cluster, from the Dutch Research Council for financial support through a grant awarded to F.L. and R.S.E (OCENW.KLEIN.498).

## Author Contributions

Conceptualisation: R.F.A.B., L.V., R.S.E., O.D.N., F.L.; Target Group Identification: R.F.A.B., L.V., R.S.E., O.D.N., F.L., J.M.F.-P., J.C.-C., S.M.-H., S.S., P.V., J.P., Y.A.F., L.S.J.-G.; Sampling: R.F.A.B., O.D.N., N.M.S.v.d.B., M.d.A.-A., C.G.-M., R.J.-M., J.C.-C., A.M., J.P., Y.A.F., L.S.J.-G., S.M.-H., S.S., P.V.; Sequencing: R.F.A.B., L.v.H., S.M.-H., J.P., Y.A.F.; Bioinformatics: L.V., L.J.; Modelling: R.F.A.B., L.V., R.S.E., O.D.N.; Writing first draft: R.F.A.B.; L.V., R.S.E., O.D.N., F.L.; Supervision: F.L., L.V., R.S.E. All co-authors contributed to editing and reviewing the manuscript.

## Data Availability

All scripts and publicly available data are available from a Zenodo repository (DOI: 10.5281/zenodo.16565207). Raw sequence data will be made available on NCBI upon acceptance.

