## Supplementary figures and images for "Trait-dependent diversification and spatio-ecological limits drive angiosperm diversity unevenness across the Canary Islands archipelago"

### Supplementary Data S5

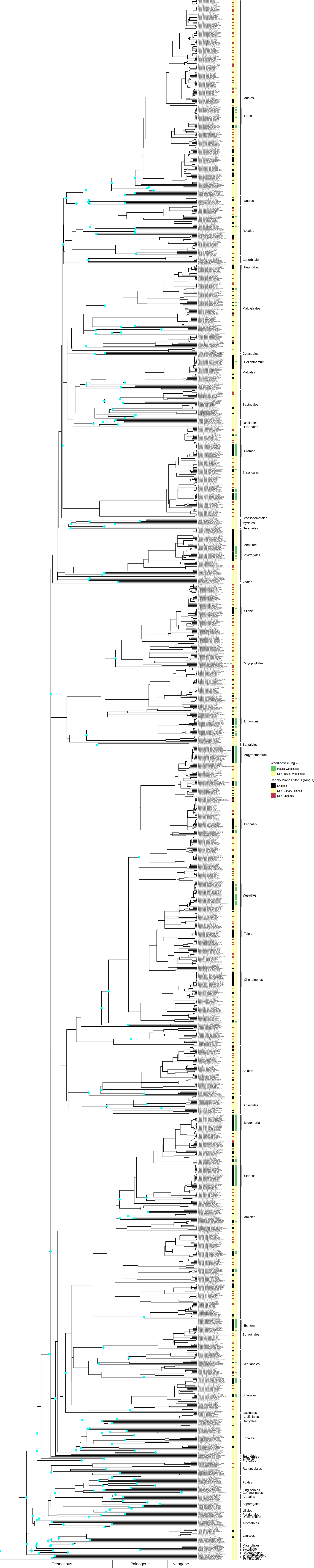

### Supplementary Figures S1 and S2

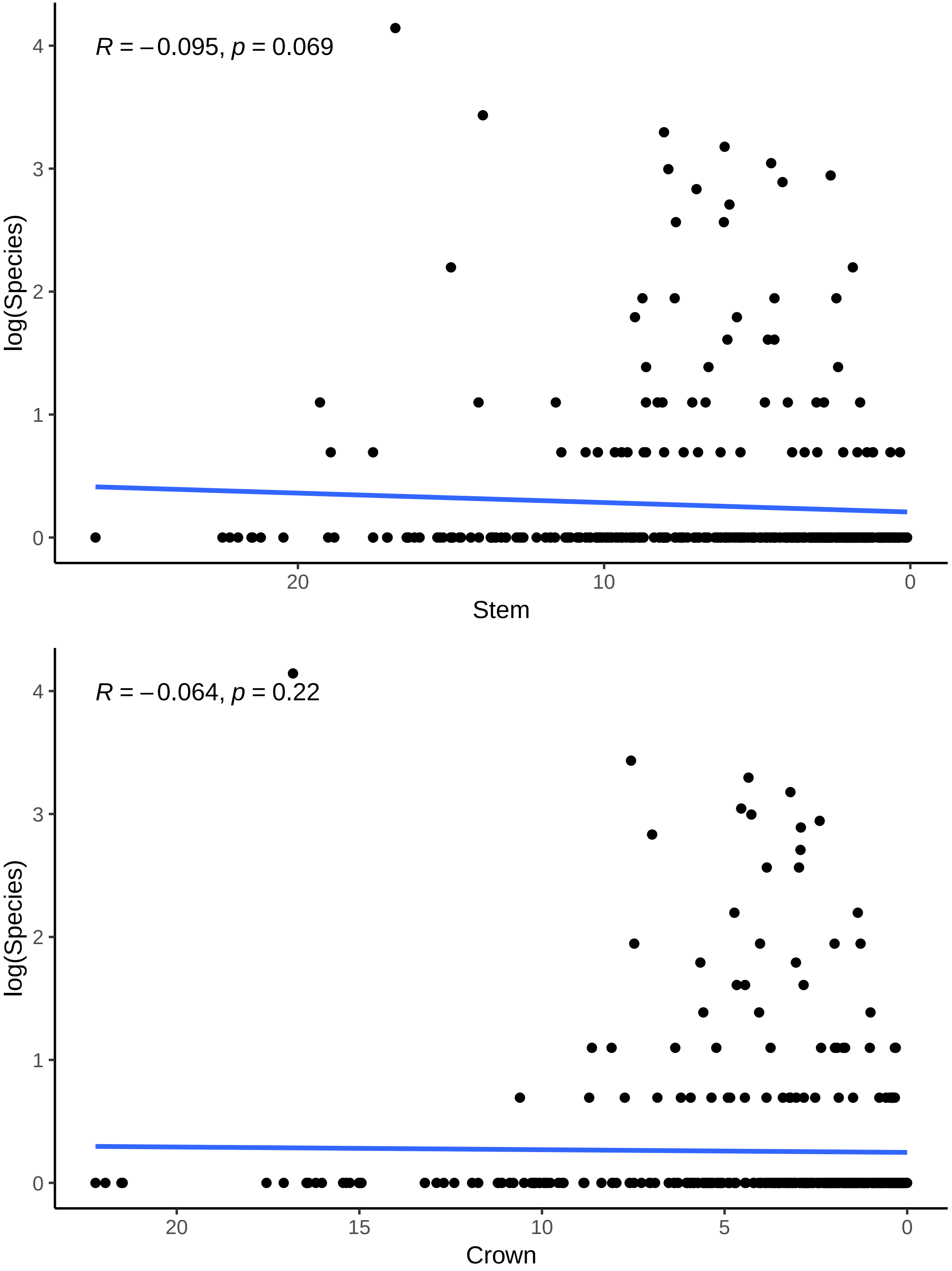

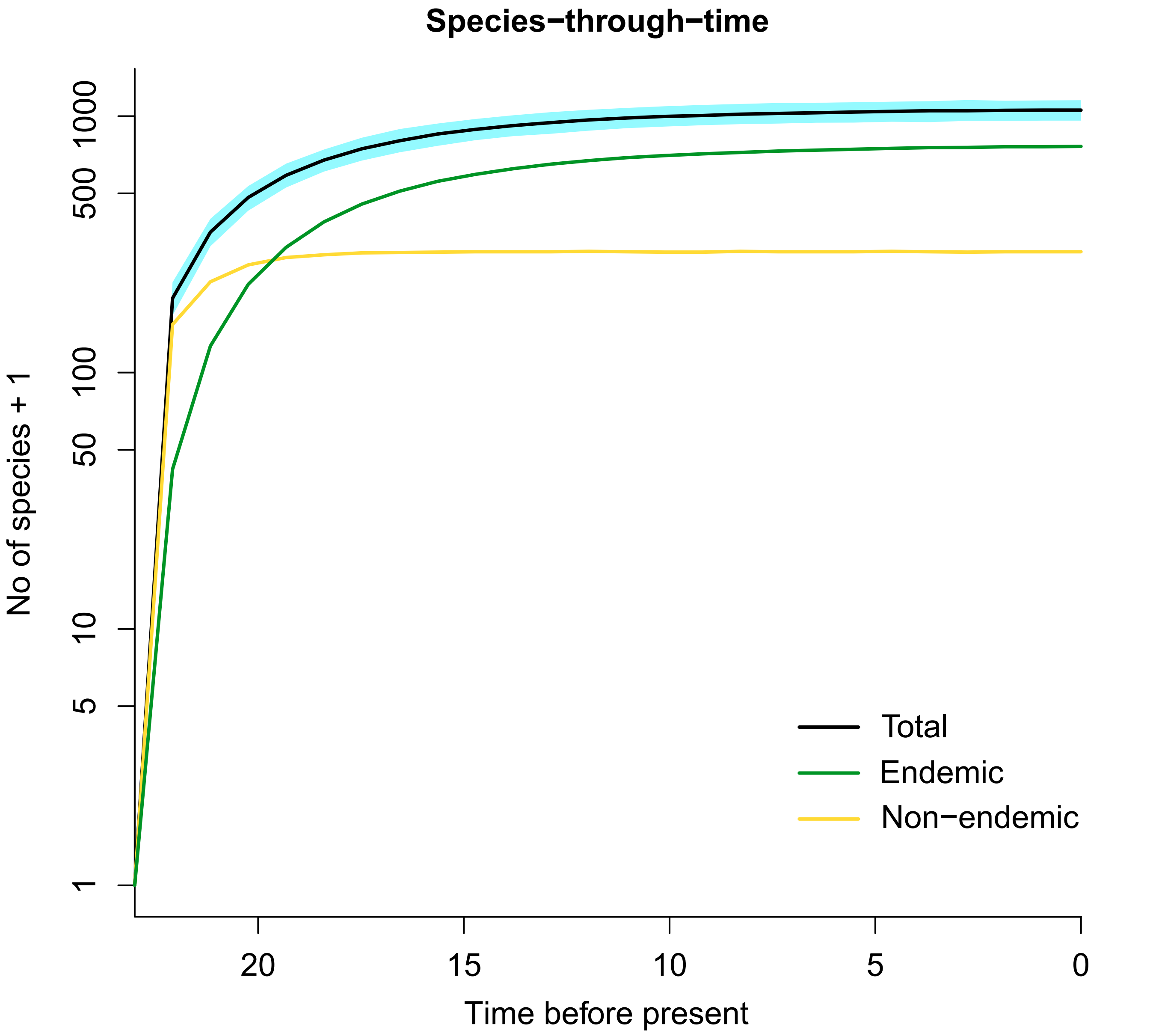
