## Supplementary Files Description for "Trait-dependent diversification and spatio-ecological limits drive angiosperm diversity unevenness across the Canary Islands archipelago"

**Supplementary Data 1**

File Name: Data_S1_Colonisation_Table_Canarian_Angiosperms

Description: Table containing the families, genera, and species considered native to the Canary Islands, with each row corresponding to a single lineage when phylogenetic data is available. The number of native species, number of colonisations (minimum, likely, and maximum), their endemicity status, whether phylogenetic data is available for the lineages, the phylogenetic data used to infer the lineages, the justification for the lineages, whether the species is potentially introduced to the Canary Islands, and whether the lineage is included in Figure 2 are recorded. Emboldened lineages indicate lineages for which no phylogenetic data was available to sufficiently resolve the number of lineages present on the Canaries. The second sheet (“In-situ_Div.”) gives the number of endemic species per genus.

**Supplementary Data 2**

File Name: Data_S2_Canary_Islands_Species_in_tree_and_MRCA

Description: The native Canary Islands species included in Figure 2, the nearest continental ancestor (using Data_S1), and the number of unique species in Figure 2. The ‘Outgroup 1’ and ‘Outgroup 2’ sheets list the native Canary Islands species in Figure 2 and our first and second choices of continental ancestors for sequencing, respectively.

**Supplementary Data 3**

File Name: Data_S3_Angiosperm_families_colonisations_and_richness

Description: The likely number of colonisations, species richness, number of genera, average number of native species per lineage, and mainland richness per native angiosperm family in the Canary Islands.

**Supplementary Data 4**

File Name: Data_S4_Binomial_Results_All_Families

Description: Results of the binomial tests to assess island disharmony per native angiosperm family in the Canary Islands, with mainland proportion, p-value, estimated binomial value, and the over-/under-representation of the family.

**Supplementary Data 5**

File Name: Data_S5_Figure_2_Vertical

Description: The vertical representation of Figure 2, with the taxonomical order, 15 largest radiations, endemicity status, and insular woodiness, as recorded in Figure 2. The blue circles at the nodes denote fossil-calibrated nodes. Fossil calibration information is given in Supplementary Data S7.

**Supplementary Data 6**

File Name: Data_S6_Final_Dated_Pruned_Clean_Con_ML_new_labels

Description: Raw phylogenetic reconstruction output from RelTime, used to construct Figure 2, with estimated divergence times given at the nodes. Tip labels correspond to the family, binomial nomenclature, sampling provenance, and sampling code as given in Supplementary Data S2.

**Supplementary Data 7**

File Name: Data_S7_Fossil_Calibrations_and_clades

Description: Nodes used for fossil calibration in RelTime, with the corresponding tip label, fossil calibration number, parent node, taxon reference, assigned node, and fossil reference. Data adapted from the ‘AngioCal’ dataset of Ramírez-Barahona et al. (2020), Nat. Ecol. Evol.

**Supplementary Data 8**

File Name: Data_S8_Pruned_Clean_Constr_CI_ML_support_values

Description: Raw phylogenetic reconstruction from RAxML-NG, with bootstrap values given at the nodes. Tip labels are as in Supplementary Data S6.

**Supplementary Data 9**

File Name: Data_S9_Woodiness Scoring Canary Islands Target Group

Description: Growth form of native Canary Islands non-monocots sampled in this study. Whether the species is included in Figure 2, the source of the growth form data, and differences with published estimates are included. The remaining insular woody species not included in this study are given in the third tab (“Difs with published estimates”).

**Supplementary Data 10**

File Name: Data_S10_Illustration_Details

Description: The source, plate number, artist, volume, issue, year of publication, link to publishing website, taxonomic information, and corresponding Canary Islands radiation of the illustration used in Figure 2.

**Supplementary Data 11**

File Name: Data_S11_Type_2_IW_Proportion

Description: Canary Island lineages with insular woody species, the corresponding tip label in Figure 2, endemicity status of the lineage, number of native species in the Canary Islands, majority growth form of the lineage, mainland relative lineage, number of species in the mainland relative lineage, number of herbaceous species in the mainland relative lineage, with the justification of the lineage, and reference used to generate the proportion of species in the differential rates (‘type 2’) other growth forms vs. insular woody DAISIE analysis.

**Supplementary Data 12**

File Name: Data_S12_Age_diversity_stem_crown_and_woodiness_status_per_lineage

Description: Species richness, stem, and crown colonisation estimates for all native Canary Islands non-monocot species, with justification of the colonisation estimate used, growth form, and tip label of the lineage in Figure 2, used in Figures 3 and 4. “Other woodiness” in this table is equivalent to either derived or ancestrally woody in Supplementary Data S9.

**Supplementary Data 13**

File Name: Data_S13_Canary_Islands_Colonisation_Time_Comparison

Description: Comparisons of our estimated stem and crown colonisation times with published estimates, including minimum and maximum confidence intervals. Adapted from Garcia-Verdugo et al. (2019), Hooft van Huysduynen et al. (2021), and Martin-Hernanz et al. (2023), including the original sources of the colonisation times used in these studies. Confidence intervals here are from Supplementary Data S14.

**Supplementary Data 14**

File Name: Data_S14_Dated_Con_ML_tree_Conf_Intervals

Description: Raw phylogenetic reconstruction in Supplementary Data S6 with the confidence intervals of estimated divergence times given at the nodes, output from RelTime. Tip labels correspond to the family, binomial nomenclature, sampling provenance, and sampling code as given in Supplementary Data S2.

**Supplementary Data 15**

File Name: Data_S15_HybPiper_Stats_All_Specimens

Description: HybPiper stats results for all sampled specimens included in Figure 2 and their corresponding tip label, with the amount of leaf material sampled, the age of the specimen, colour of the leaf material, initial DNA extraction quantity (in ng/ul), DNA quantity after library preparation, the source collection of the material, and the herbarium code of the sampled specimen.

**Supplementary Data 16**

File Name: Data_S16_MO_Paralogy_resolved_astral_tree

Description: Paralog filtered phylogenetic reconstruction using 111 orthologous groups output from ParaGone, constructed using ASTRAL-IV. Tip labels are as in Supplementary Data S6.

**Supplementary Data 17**

File Name: Data_S17_Updated_DAISIE_Table_Full_Stem_Crown

Description: Input table used to run DAISIE for both the stem and stem+crown datasets, with the name of the clade (as in Figure 2), the endemicity status, number of missing species from our phylogenetic reconstruction, branching times, and the source of the branching times used. Adapted to create Supplementary Data S12 and Figures 3 and 4.
